# MdLRR-RLK1-MdATG3 module enhances the resistance of apples to abiotic stress via autophagy

**DOI:** 10.1101/2024.09.20.614134

**Authors:** Wenjun Chen, Wei Guo, Chao Zhang, Yi Zhao, Yingying Lei, Cui Chen, Ziwen Wei, Hongyan Dai

## Abstract

LRR-RLKs play a key role in plant responses to stress, although their physiological functions under abiotic stress are not yet fully understood. *MdLRR-RLK1* was found to regulate plant growth and development and to increase tolerance to salt and drought stress. MdLRR-RLK1 interacts with MdATG3, which ubiquitinates and degrades MdLRR-RLK1. MdATG3 enhances salt and drought tolerance through autophagy, interacting with MdATG8F/I-like in apple. These findings reveal the interaction between MdLRR-RLK1 and MdATG3, suggesting mechanisms that regulate apple growth and resistance to abiotic stress.

## Introduction

Apples are highly cultivated and economically valuable fruits worldwide and are cherished by people for their high nutritional content (Wang et al., 2020). The growth of apples is often restricted by environmental factors such as water availability and salinity, especially in the face of climate change (Xiong et al., 2001; Liu et al., 2024). Freshwater, which accounts for less than 3% of the Earth’s total supply, is heavily consumed by agriculture, with a current usage rate of 70%. Despite environmental changes, freshwater consumption continues to rise. Furthermore, the apple cultivation industry, which is primarily situated in Mediterranean and subtropical regions, is negatively impacted by ecological disturbances, and excessive water consumption in agriculture remains an urgent issue to resolve (Diffenbaugh et al., 2005; White et al., 2006; Stocker, 2013). Soil salinity is another influential factor affecting plant productivity, with data suggesting that approximately 50% of arable land will face salinization issues in the near future (Zhang et al., 2023). As a result, understanding the physiological changes and underlying molecular mechanisms of the response of apples to abiotic stress is crucial for the sustainable development of the apple cultivation industry and overall socioeconomic harmony. Consequently, there is an immediate need to cultivate apple varieties that are tolerant to drought and salt stresses.

Plants have developed numerous receptor-like kinases (RLKs), which are classified on the basis of their kinase and cellular domains (Gish and Clark, 2011; Osakabe et al., 2013). The most abundant subcategory of RLKs is the leucine repeat sequence (*LRR-RLK*) group. *LRR-RLKs* not only contribute to plant development and various pathways but also play a distinctive role in plant responses to stress and challenging conditions (Dievart and Clark, 2004; Li et al., 2018). Research has established the critical involvement of *LRR-RLKs* in abiotic stress. For example, Park et al. (2014) reported that RLK, *OsGIRL1*, which contains leucine repeat sequences, is significantly induced under abiotic stressors such as salt, drought, salicylic acid (SA), and abscisic acid (ABA). *OsGIRL1* promoted growth and development upon heterologous overexpression in Arabidopsis. Additionally, Liu et al. (2020) demonstrated that the LRR-RLK protein AtHSL3 can regulate drought tolerance and sensitivity to ABA in Arabidopsis. Moreover, Lin et al. (2020) reported that the overexpression of *OsSTLK*, an LRR-RLK protein rich in leucine repeat sequences, can increase salt stress tolerance, substantially reducing malondialdehyde (MDA) content and mitigating reactive oxygen species (ROS) accumulation. Furthermore, the ATTED database (https://atted.jp/hclust/) suggests potential interaction mechanisms among *LRR-RLKs*; however, further experimental verification is essential (Cui et al., 2018).

Autophagy is a highly conserved process that takes place in eukaryotic cells, allowing for the degradation or recovery of unnecessary cellular components and the maintenance of cellular homeostasis under both normal and stressful conditions (Liu and Bassham, 2012; Marshall and Vierstra, 2018; Su et al., 2020). Autophagy has also been shown to play a significant role in drought and salt stress responses. Previous studies have demonstrated that several autophagy-related genes (*ATGs*), such as *CaATG2* in *Capsicum annuum* L., *SiATG8a* in *Setaria italica* var. germanica (Mill.) Schred., *HvATG6* in *Hordeum vulgare* L., and *MdATG3* and *MdATG18a* in *Malus domestica* Borkh (Wada et al., 2015; Li et al., 2016; Zhai et al., 2016; Wang et al., 2017; Zeng et al., 2017). Additionally, Wang et al. (2015) reported that silencing *SlATG10* and *SlATG18f* in tomatoes (*Solanum lycopersicum* L.) led to the inhibition of autophagosome formation and reduced drought tolerance. Similarly, Sun et al. (2018) and Li et al. (2020) reported that *MdATG18a* and *MtCAS31* could impact plant drought tolerance by affecting autophagy. Furthermore, the AtPI3K complex in Arabidopsis was found to increase salt tolerance by promoting AtPIP2 (Huo et al., 2020), whereas the overexpression of *MdATG10* in apples increased root autophagy activity and improved apple salt stress tolerance (Zhang et al., 2021).

Previous studies have shown that heterologous overexpression of *MdATG3* in Arabidopsis can increase tolerance to abiotic stress, and the same result was observed for *ZmATG3* overexpression in Arabidopsis (Wang et al., 2017; Liu et al., 2024). Moreover, ATG3 functions as an E2 ubiquitin-binding enzyme, catalyzing the binding of proteins and lipids (Hanada et al., 2009). ATG3 is a critical component of autophagy and plays a role in the ubiquitination process. The binding of PE to the C-terminal Gly of ATG8 results in the formation of an ATG8-PE conjugate, further promoting autophagy (Ichimura et al., 2000; Kirisako et al., 2000; Yamaguchi et al., 2012; Huang et al., 2023). In Arabidopsis, ATG8 has been shown to increase autophagy by regulating its binding to ATG8 (Guan and Xue, 2022), indicating that ATG3 functions similarly to an E2 ubiquitin-binding enzyme in the regulation of autophagy.

A previous study revealed that the overexpression of *CpLRR-RLK1* improved tolerance to *Pst* DC3000 in *Arabidopsis thaliana.* Owing to the high sequence similarity between the CDSs of *CpLRR-RLK1* and *MdLRR-RLK1*, along with existing research on *LRR-RLKs*, *MdLRR-RLK1* was selected as the main research focus. Our findings indicate that MdLRR-RLK1 enhances GL-3 tolerance to abiotic stress and is degraded by MdATG3 through phosphorylation followed by ubiquitination. These results provide a theoretical basis for developing apple varieties with improved tolerance to abiotic stress.

## Results

### Characterizations of *MdLRR-RLK1* expression

To examine the phylogenetic relationship of *MdLRR-RLK1*(Mdg_15A013100), a comparison was made with the sequences of the LRR-RLK family genes from Arabidopsis. The analysis revealed that *MdLRR-RLK1* is closely related to the Arabidopsis SERK gene family (Supplementary Fig. S1A). Structural domain comparisons between *MdLRR-RLK1* and Arabidopsis SERKs revealed that *MdLRR-RLK1* contains an LRR domain, a transmembrane (TM) region, and an S-TKc domain (Supplementary Fig. S1B). The expression of *MdLRR-RLK1* in various GL-3 tissues was highest in the roots, suggesting its involvement in stress responses (Supplementary Fig. S1F). To further explore its time‒response pattern, GL-3 plants were treated with mannitol and NaCl, and *MdLRR-RLK1* expression was significantly induced at 72 h under both treatments (Supplementary Fig. S1C, D). Subcellular localization experiments confirmed that *MdLRR-RLK1* is localized to the cell membrane (Supplementary Fig. S1E).

### *MdLRR-RLK1* could regulate plant growth and development in apple

To further investigate the function of *MdLRR-RLK1*, four *MdLRR-RLK1*-overexpressing lines and sixteen *MdLRR-RLK1* RNA interference lines were obtained. Two overexpressed lines, named OE-1 and OE-2, along with two RNA interference lines, named RNAi-5 and RNAi-15, were selected for subsequent experiments (Fig. 1A, B). After these transgenic lines were cultivated in the culture room for three months, we evaluated various morphological indicators, including plant height, leaf length, leaf width, number of leaves, number of internodes, and internode length. The results revealed that plant height significantly increased by 17.0%-20.2% in OE-1 and OE-2, whereas compared with that in GL-3, the height in both RNAi-5 and RNAi-15 significantly decreased by 23.7%-25.6% (Fig. 1C). Similarly, OE-1 and OE-2 presented significant increases of 11.8%-21.8% in terms of leaf length, leaf width, number of leaves, number of internodes, and internode length, whereas RNAi-5 and RNAi-15 presented significant decreases in these morphological indicators (Fig. 1D-H). During cultivation, the root number of OE-1 and OE-2 was greater than that of GL-3, RNAi-5, and RNAi-15, although no significant difference was detected among these lines, except for RNAi-5, which presented a shorter root length (Fig. 1I, J). Overall, these findings indicate that *MdLRR-RLK1* may play a role in the regulation of plant growth and development.

**Fig. 1.**
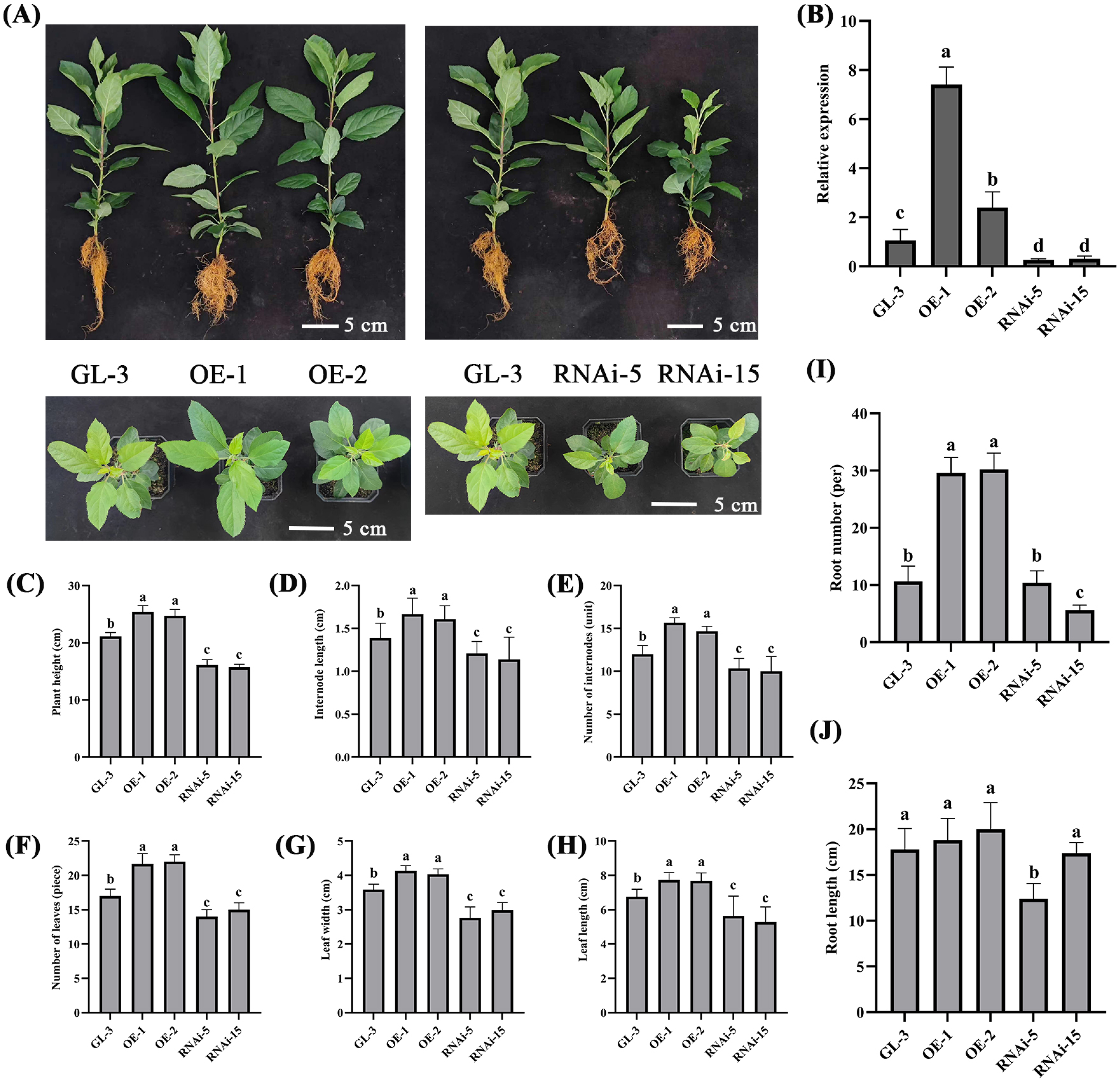
Phenotypic characterization of the MdLRR-RLKl transgenic lines. (A) Phenotypes of the MdLRR-RLKl overexpression and RNA interference lines. (B) RT-qPCR analysis of MdLRR-RLKl expression in the overexpression and RNA interference lines. (C) Comparisons of plant height among GL-3, MdLRR-RLKl-overexpressing, and RNA interference lines. (D) Comparison of the internode lengths of the GL-3, MdLRR-RLKl-overexpressing, and RNA interference lines. (E) Comparison of intemode number among GL-3, MdLRR-RLKl-overexpressing, and RNA interference lines. (F) Comparative leaf number counts of the GL-3, MdLRR-RLKl-overexpressing, and RNA interference lines. (G) Comparative leaf width measurements of GL-3, MdLRR-RLKl-overexpressing, and RNA interference lines. (H) Comparative leaf length measurements of the GL-3, MdLRR-RLKl-overexpressing, and RNA interference lines. (I) Root numbers of the GL-3, MdLRR-RLKl-overexpressing, and RNA interference lines. The error bars represent the standard deviations (n ≥ 3, P < 0.05). (J) Root length of GL-3, MdLRR-RLKl-overexpressing, and RNA interference lines. The error bars represent the standard deviation (n ≥ 3, P < 0.05). OE refers to the overexpression lines of MdLRR-RLKl, and RNAi refers to the RNA interference lines of MdLRR-RLKl. The scale bar represents 5 cm. The vertical bars indicate the standard deviation. Different letters denote significant differences compared with GL-3 (n ≥ 3, P < 0.05).

### Overexpressing *MdLRR-RLK1* increased tolerance to salt and drought stress in apple

Given the significant induction of *MdLRR-RLK1* expression by salt stress in GL-3, it was hypothesized that *MdLRR-RLK1* may be involved in the salt stress response. To investigate this phenomenon, OE-1, OE-2, RNAi-5, RNAi-15, and GL-3 were subjected to 200 mM NaCl treatment. After 15 days, RNAi-5 and RNAi-15 resulted in leaf rolling and wilting symptoms. By day 30, RNAi-5, RNAi-15, and GL-3 resulted in more severe leaf drooping and chlorosis than OE-1 and OE-2 did. After 60 days, RNAi-5, RNAi-15, and GL-3 resulted in signs of mortality, whereas OE-1 and OE-2 remained viable (Fig. 2A). Enzyme activity analysis revealed that POD, CAT, and SOD activities were significantly greater in OE-1 and OE-2 than in GL-3 under salt stress, whereas the activities of RNAi-5 and RNAi-15 were significantly lower than those of GL-3 (Fig. 2B-D). Furthermore, the MDA content indicated that OE-1 and OE-2 resulted in relatively less leaf damage than did GL-3, RNAi-5, and RNAi-15 under salt stress (Fig. 2E). Analysis of key genes in the salt overly sensitive (SOS) pathway of the *MdLRR-RLK1* transgenic lines under NaCl treatment revealed that the expression levels of *MdSOS1*, *MdSOS2*, and *MdSOS3* were all increased, with greater expression in the overexpression lines than in the GL-3 lines, whereas the RNAi lines presented the opposite trend (Fig. 2F-H). Similarly, OE-1 and OE-2 presented increased drought tolerance (Fig. 3A). Furthermore, the activities of POD, CAT, and SOD in OE-1 and OE-2 were significantly greater than those in GL-3, RNAi-5, and RNAi-15 under drought treatment. Analysis of the MDA contents indicated that OE-1 and OE-2 resulted in less leaf damage than did GL-3, RNAi-5, and RNAi-15 (Fig. 3B-D). Additionally, the expression of the wax synthesis-related genes *MdMYB16*, *MdMYB94*, and *MdSPS6* was significantly greater in the overexpression lines than in GL-3 (Fig. 3F-H). These findings suggest that *MdLRR-RLK1* exhibits increased antioxidant enzyme activity and increased tolerance to salt and drought stress.

**Fig. 2.**
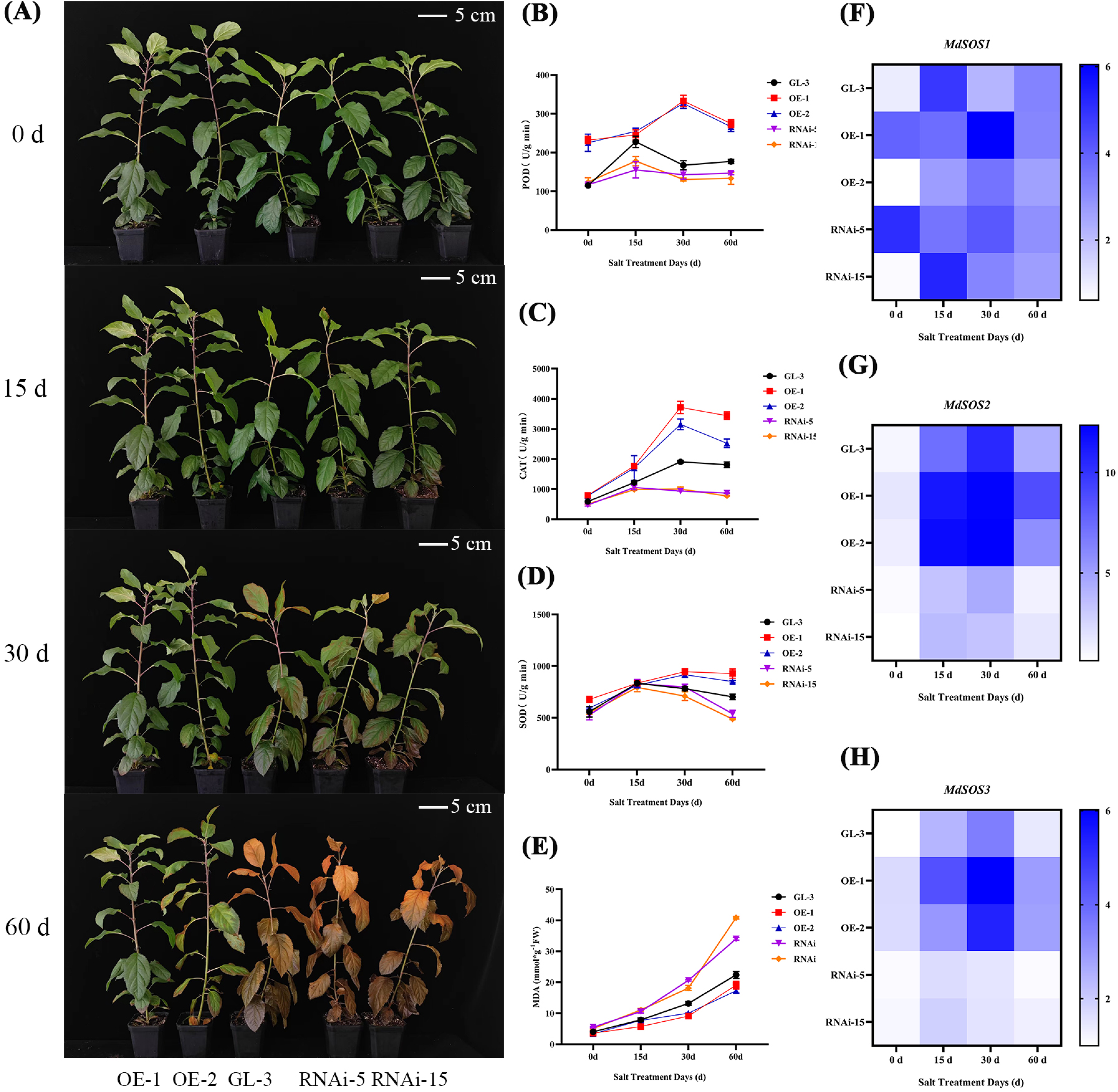
Phenotypes of the MdLRR-RLKl transgenic lines and GL-3 under salt stress. (A) Phenotypes of the MdLRR-RLKl transgenic lines and GL-3 plants under salt treatment at 0, 15, 30, and 60 days. (B) Line graph of the POD activity of the MdLRR-RLKl transgenic lines and GL-3 plants under salt treatment at 0, 15, 30, and 60 days. (C) Line graph of the CAT activity of the MdLRR-RLKl transgenic lines and GL-3 plants under salt treatment at 0, 15, 30, and 60 days. (D) Line graph of the SOD activity of the MdLRR-RLKl transgenic lines and GL-3 plants under salt treatment at 0, 15, 30, and 60 days. (E) Line graph of the MDA content of the MdLRR-RLKl transgenic lines and GL-3 plants under salt treatment at 0, 15, 30, and 60 days. (F) MdSOSl expression levels in MdLRR-RLKl transgenic lines and GL-3 plants under salt treatment at 0, 15, 30, and 60 days. (G) MdSOS2 expression levels in MdLRR-RLKl transgenic lines and GL-3 plants under salt treatment at 0, 15, 30, and 60 days. (H) MdSOS3 expression levels in MdLRR-RLKl transgenic lines and GL-3 plants under salt treatment at 0, 15, 30, and 60 days.

**Fig. 3.**
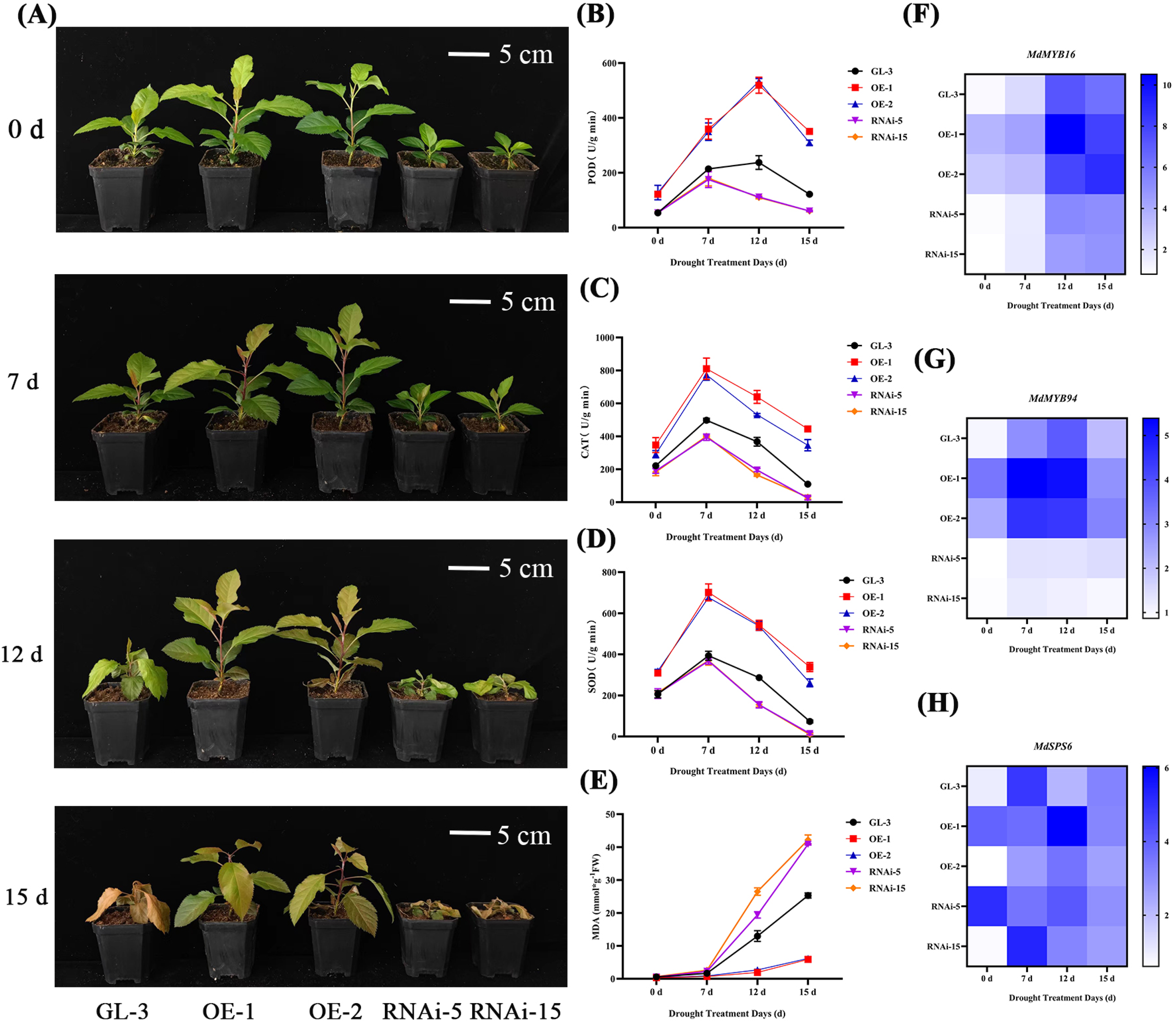
Phenotypes of the MdLRR-RLKl transgenic lines and GL-3 plants under drought treatment_(A) Phenotypes of the MdLRR-RLKl transgenic lines and GL-3 plants under drought treatment at 0, 7, 12, and 15 days_(B) Line graph of the POD activity of the MdLRR-RLKl transgenic lines and GL-3 plants under drought treatment at 0, 7, 12, and 15 days_(C) Line graph of the CAT contents of the MdLRR-RLKl transgenic lines and GL-3 plants under drought treatment at 0, 7, 12, and 15 days. (D) Line graph of the SOD activity of the MdLRR-RLKl transgenic lines and GL-3 plants under drought conditions at 0, 7, 12, and 15 days. (E) Line graph of the MDA contents of the MdLRR-RLKl transgenic lines and GL-3 plants under drought treatment at 0, 7, 12, and 15 days. (F) MdMYB16 expression levels in MdLRR-RLKl transgenic lines and GL-3 plants under drought conditions at 0, 7, 12, and 15 days. (G) MdMYB94 expression levels in MdLRR-RLKl transgenic lines and GL-3 plants under drought conditions at 0, 7, 12, and 15 days. (H) MdSPS6 expression levels in the MdLRR-RLKl transgenic lines and GL-3 plants under drought treatment at 0, 7, 12, and 15 days.

### *MdLRR-RLK1* transcriptome analysis

For a closer examination of the molecular mechanisms associated with *MdLRR-RLK1*, this study conducted transcriptome sequencing and KEGG pathway analysis on GL-3 and the line with the highest expression level, OE-1. The findings revealed significant upregulation of genes related to macrophages, POD, and plant hormones (Supplementary Fig. S2A). Autophagy pathway gene analysis revealed the upregulation of most autophagy genes in the overexpression lines (Supplementary Fig. S2B). Heatmap analysis of peroxisome pathway genes revealed increased expression of POD-related genes in the overexpression materials (Supplementary Fig. S2C). These results suggest that the overexpression of LRR-RLK1 may increase stress resistance.

### MdLRR-RLK1 interacted with MdATG3

To investigate the mechanism of *MdLRR-*RLK1 resistance to drought and salt stress, the *MdLRR-RLK*-BT3-STE vector was constructed for yeast library screening, leading to the identification of *MdATG3*(Mdg_02B018380) as the interacting partner. The interaction between *MdLRR-RLK1* and *MdATG3* was further validated via yeast two-hybrid (Y2H) assays with the *MdLRR-RLK*-BT3-STE and *MdATG3*-PR3-N vectors, confirming the interaction in yeast cells (NMY51) (Fig. 4A). Additionally, BiFC assays were performed to verify the interaction in plants. The results revealed that *MdLRR-RLK1*-YFP^n^ and *MdATG3*-YFP^c^ interacted in the cell membrane of tobacco cells, whereas negative controls (*MdLRR-RLK1*-YFPn/YFP^c^ and YFPn/*MdATG3-*YFP^c^) did not produce YFP signals (Fig. 4C). Pull-down assays also demonstrated that *MdLRR-RLK1*-His could interact with *MdATG3*-GST (Fig. 4D). Additionally, split luciferase complementation (SLCA) assays in which *MdLRR-RLK1* was fused with N-terminal luciferase (*MdLRR-RLK1*-nLUC) and *MdATG3* was fused with C-terminal LUC (*MdATG3*-cLUC) revealed increased luciferase activity in tobacco leaves, confirming the interaction. The negative controls (*MdLRR-RLK1*-nLUC/cLUC, nLUC/*MdATG3*-cLUC, and nLUC/cLUC) presented lower luciferase activity (Fig. 4B). These results indicated that *MdLRR-RLK1* could interact with *MdATG3* both *in vivo* and *in vitro*.

**Fig. 4.**
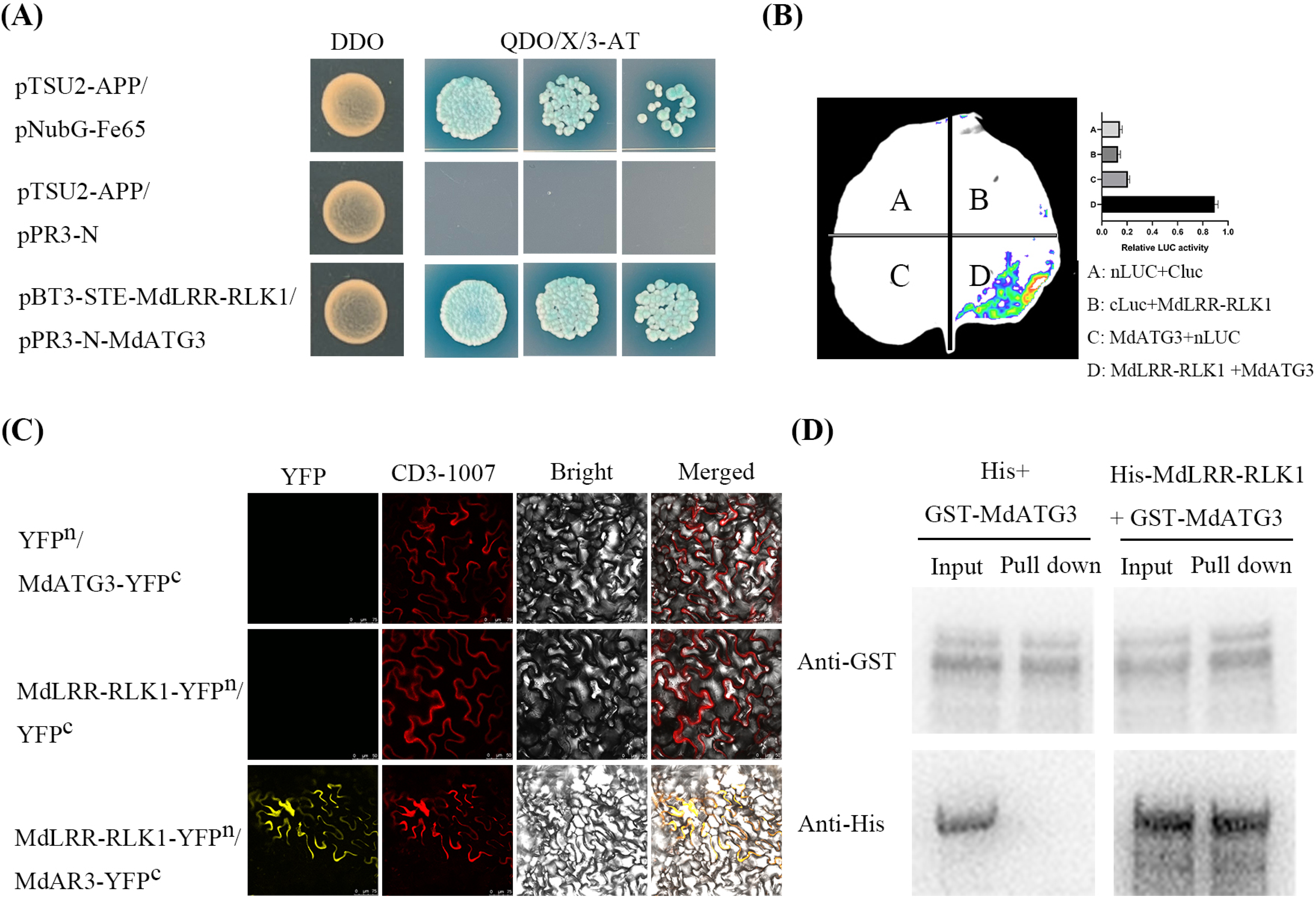
MdLRR-RLKl interacts with MdATG3 both in vitro and in vivo. (A) Yeast 2 hybrid validation. PBT3-STE-MdLRR-RLKl, pPR3-N-MdATG3, and control plasmids were cotransformed into NMY51 competent yeast, which were subsequently cultured on media for 3-5 days after transformation. (B) Luciferase imaging assay. MdLRR-RLKl-nLUC, MdATG3-cLUC, and the control plasmid were injected into tobacco leaves via Agrobacterium tumefaciens. What method (reference or reagent kit) should be used to determine LUC content. (C) BiFC assay analysis. The MdLRR-RLKl-YFPn, MdATG3-YFPc, and control plasmids were subsequently injected into tobacco leaves via Agrobacterium tumefaciens. The control group had scales of75 µm and 50 µm, whereas the experimental group had a scale of75 µm. (D) Pull-down test analysis. Escherichia coli containing MdLRR-RLKl-His, MdATG3-GST, and empty plasmids were induced to obtain the corresponding proteins, which were detected via SDS-PAGE.

### MdATG3 can ubiquitinate and degrade MdLRR-RLK1

This study utilized phosphatase inhibitors during the cleavage of the MdLRR-RLK1 and MdATG3 proteins and subsequently conducted WB detection of the extracted proteins. Our findings revealed that the bands corresponding to the MdLRR-RLK1 protein extracted with phosphate inhibitors appeared at a greater position than those corresponding to the protein extracted without phosphatase inhibitors (Supplementary Fig. S3A). Then, analyzed the MdLRR-RLKl protein extracted with phosphate inhibitors by liquid chromatography-tandem mass spectrometry (LC-MS/MS). Only one tyrosine residue (Y347) on MdLRR-RLK1 protein was phosphorylated (Supplementary Fig. S3B). This finding suggests that phosphorylation may occur with MdLRR-RLK1. To further investigate this phenomenon, *in vitro* phosphorylation assays were performed using MdLRR-RLK1 extracted with and without phosphatase inhibitors to assess its potential to phosphorylate MdATG3. The results indicated that MdLRR-RLK1 did not phosphorylate MdATG3 *in vitro* (Fig. 5A, B). Since MdATG3 is an E2 ubiquitin-conjugating enzyme, *in vitro* ubiquitination experiments were conducted to determine whether MdATG3 can ubiquitinate and degrade MdLRR-RLK1. The results showed that MdATG3 could ubiquitinate and degrade phosphorylated MdLRR-RLK1 but not the nonphosphorylated MdLRR-RLK1 protein (Fig. 5C, D). These findings indicate that the degradation of MdLRR-RLK1 by MdATG3 might depend on its phosphorylation.

**Fig. 5.**
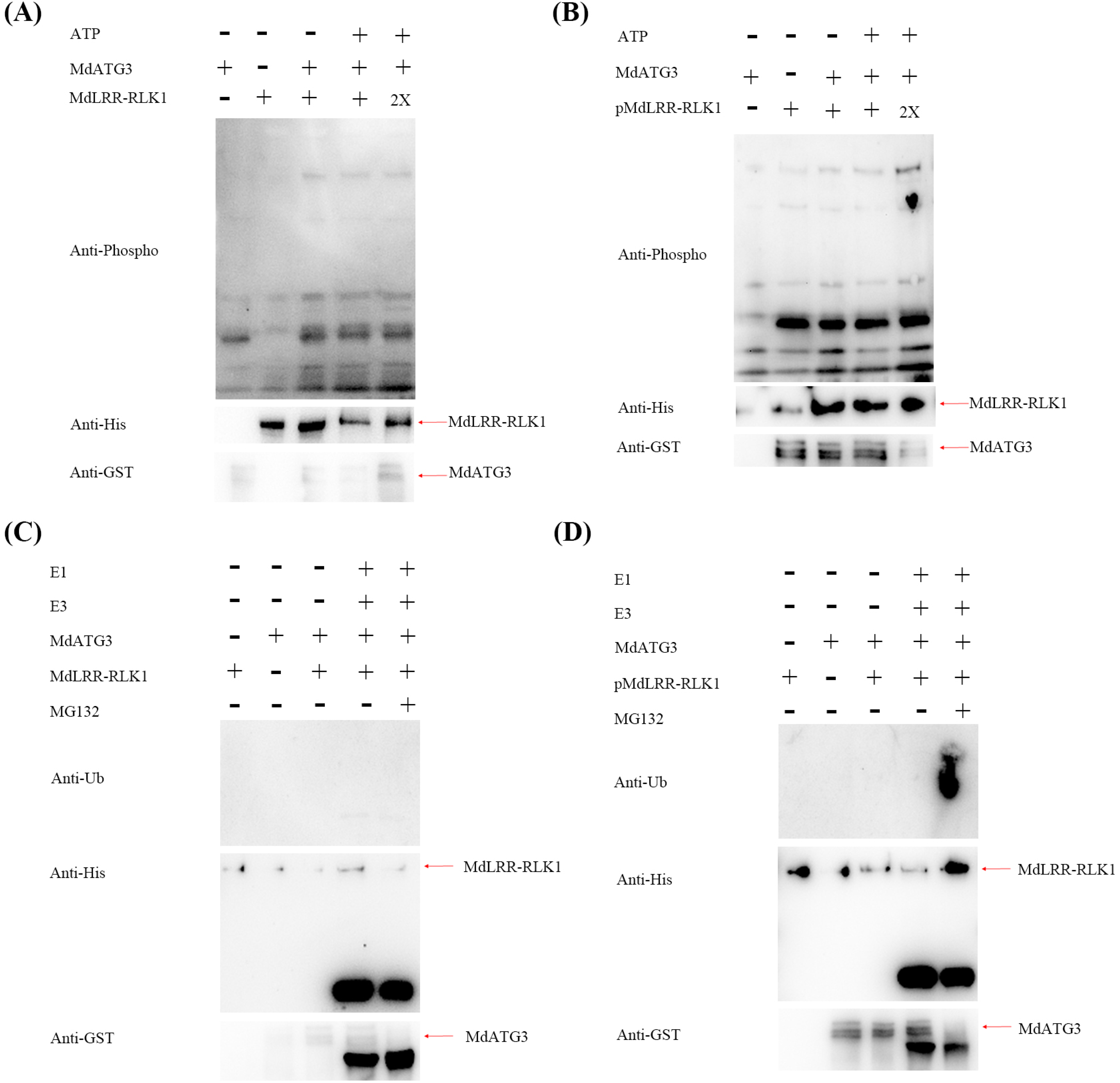
MdATG3 t1biqt1itination degrades the MdLRR-RLKl phosphorylated protein. (A) MdLRR-RLKl does not phosphorylate MdATG3 in vitro. Escherichia coli containing MdLRR-RLKl-His and MdATG3-GST was indt1ced, separated by SDS gel electrophoresis and detected with an anti-phospho serine/treonene mixtt1re. (B) pMdLRR-RLKl does not phosphorylate MdATG3 in vitro. pMdLRR-RLKl (protein of MdLRR-RLKl t1nder PIC treatment) and MdATG3-GST were separated via SDS gel electrophoresis and detected with an anti-phospho serine/treonene antibody. (C) MdATG3 does not t1biqt1itinate or degrade MdLRR-RLKl. Escherichia coli containing MdLRR-RLKl-His and MdATG3-GST was indt1ced, separated by SDS gel electrophoresis and detected with an anti-Ub antibody. (D) MdATG3 t1biqt1itinates and degrades pMdLRR-RLKl. pMdLRR-RLKl (protein of MdLRR-RLKl t1nder PIC treatment) and MdATG3-GST were separated via SDS gel electrophoresis and detected with an anti-Ub antibody.

### *MdATG3* could regulate plant growth and development in apple

Since MdLRR-RLK1 interacts with MdATG3, we conducted a structural analysis of *MdATG3* and marked the positions of the exons (Supplementary Fig. S4A). Through systematic evolutionary analysis, we discovered that MdATG3 is most closely related to PuATG3 in *Pyrus × bretschneideri* (Supplementary Fig. S4B). To further investigate this phenomenon, we examined the expression pattern of *MdATG3* and observed the highest expression level in the fruit (Supplementary Fig. S4C). Additionally, subcellular localization experiments confirmed that MdATG3 is localized on the plant cell membrane (Supplementary Fig. S4F). For quantitative analysis, we obtained 6 lines with *MdATG3* overexpression and selected *MdATG3*-1 and *MdATG3*-2 for subsequent experiments (Supplementary Fig. S4D). After these transgenic lines were cultured under artificial light for two months (Supplementary Fig. S4E), we evaluated various morphological traits, such as plant height, leaf length, leaf width, leaf number, internode number, and internode length (Supplementary Fig. S4G-L). The results demonstrated a significant increase in plant height of 35.1% and 46.0% for OE-1 and OE-2, respectively, compared with that of GL-3. Similarly, OE-1 and OE-2 presented significant increases ranging from 21.4% to 46.4% in terms of leaf length, leaf width, leaf number, internode number, and internode length. These findings suggest that *MdATG3* may play a role in regulating plant growth and development.

### *MdATG3* enhanced tolerance to salt and drought stress in apple

A significant increase in *MdATG3* expression was observed when *MdLRR-RLK1* was subjected to salt stress. To explore this further, salt treatment was applied to ATG3-1, ATG3-2, and GL-3 via 200 mM NaCl. After 3 days, GL-3 presented slight wilting, which became severe by day 7. By day 15, GL-3 had died, whereas ATG3-1 and ATG3-2 exhibited only mild wilting (Fig. 6A). Analysis of the activity of antioxidant enzymes, including POD, CAT, SOD, and MDA, revealed increased POD, CAT, and SOD activities and decreased MDA levels in ATG3-1 and ATG3-2, indicating reduced damage (Fig. 6B-E). These findings suggest that *MdATG3* enhances autophagy activity. Additionally, key genes in the SOS pathway, such as *MdSOS1*, *MdSOS2*, and *MdSOS3*, were induced by salt stress, with higher expression levels of ATG3-1 and ATG3-2 than of GL-3 (Fig. 6F-H). A drought stress experiment was also conducted in which ATG3-1, ATG3-2, and GL-3 were subjected to water shortage. After 9 days, GL-3 exhibited leaf death, whereas ATG3-1 and ATG3-2 showed only slight dehydration (Fig. 7A). Further analysis of POD, CAT, SOD, and MDA levels confirmed stronger drought tolerance in ATG3-1 and ATG3-2 (Fig. 7B-E). Compared with those of GL-3, the expression levels of the wax synthesis-related genes *MdMYB16* and *MdMYB94* were greater in ATG3-1 and ATG3-2 under drought stress (Fig. 7F, G). These findings indicate that *MdATG3* enhances antioxidant enzyme activity and improves salt and drought tolerance in apple.

**Fig. 6.**
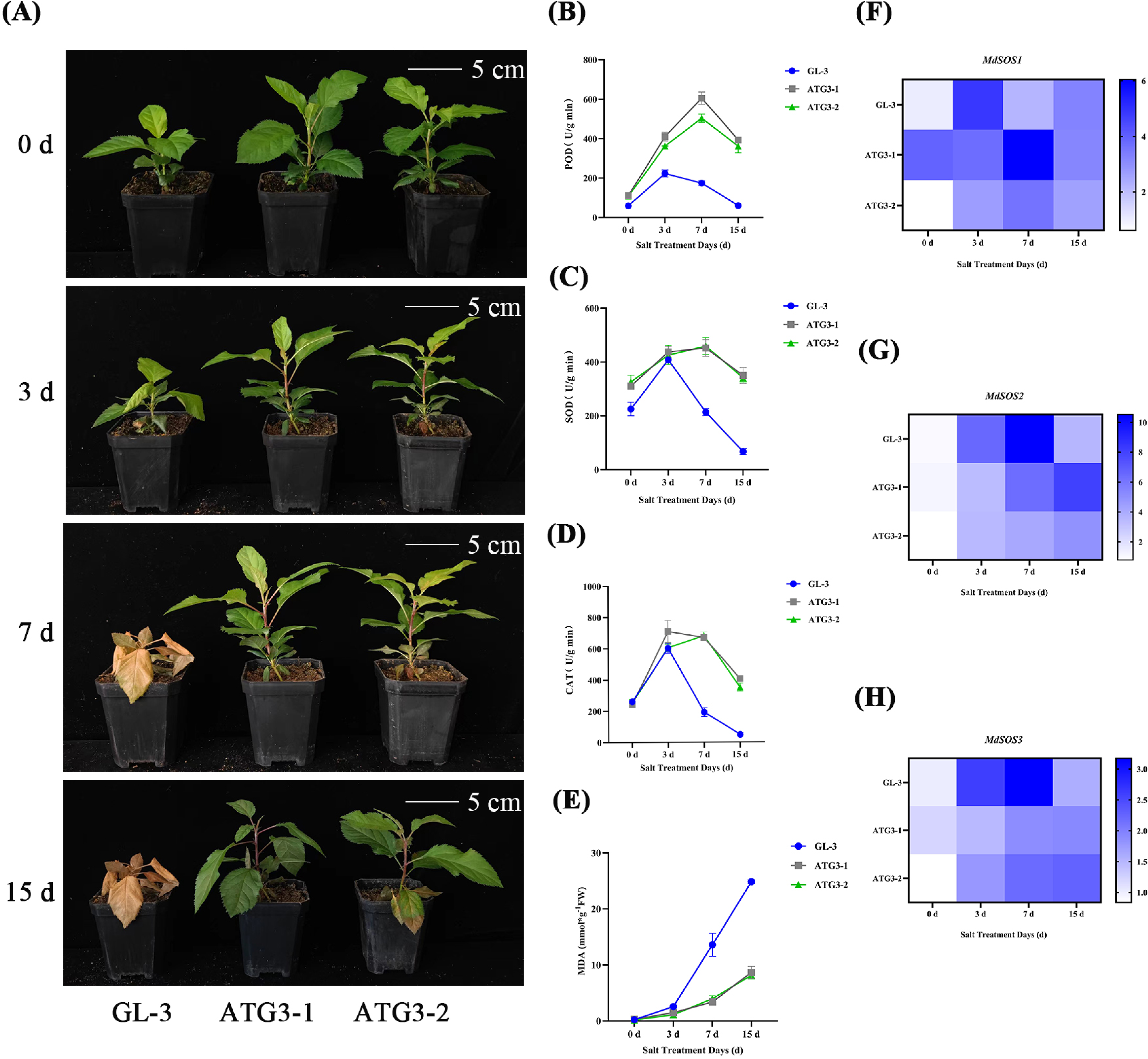
Phenotypic responses of the MdATG3-overexpressing lines and GL-3 to salt stress. (A) Phenotypes of the MdATG3-overexpressing lines and GL-3 at 0, 3, 7, and 15 days of salt treatment. (B) POD activity in the MdATG3-overexpressing lines and GL-3 at 0, 3, 7, and 15 days of salt treatment. (C) CAT content in the MdATG3-overexpressing lines and GL-3 at 0, 3, 7, and 15 days of salt treatment. (D) SOD activity in the MdATG3-overexpressing lines and GL-3 at 0, 3, 7, and 15 days of salt treatment. (E) MDA content in the MdATG3-overexpressing lines and GL-3 at 0, 3, 7, and 15 days of salt treatment. (F) Expression levels of MdSOSl in the MdATG3-overexpressing lines and GL-3 at 0, 3, 7, and 15 days of salt treatment. (G) Expression levels of MdSOS2 in the MdATG3-overexpressing lines and GL-3 at 0, 3, 7, and 15 days of salt treatment. (H) Expression levels of MdSOS3 in the MdATG3-overexpressing lines and GL-3 at 0, 3, 7, and 15 days of salt treatment.

**Fig. 7.**
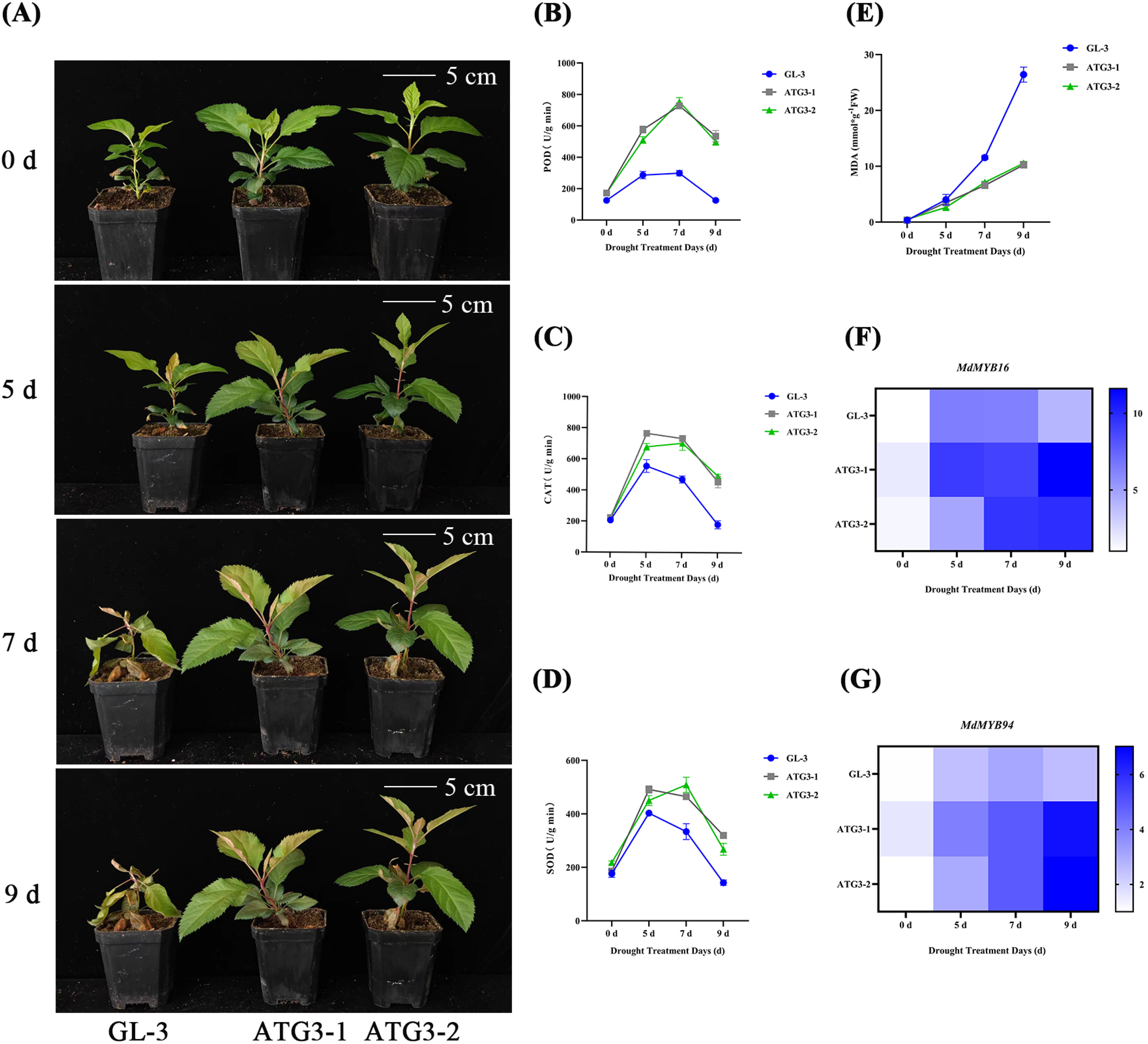
Phenotypes of the MdATG3-overexpressing lines and GL-3 under drought treatment. (A) Phenotypes of the MdATG3-overexpressing lines and GL-3 plants under drought treatment at 0, 5, 7, and 9 days. (B) Line graph of POD activity in the MdATG3-overexpressing lines and GL-3 plants under drought conditions at 0, 5, 7, and 9 days. (C) Line graph of the CAT content in the MdATG3-overexpressing lines and GL-3 plants under drought conditions at 0, 5, 7, and 9 days. (D) Line graph of the SOD activity in the MdATG3-overexpressing lines and GL-3 plants under drought conditions at 0, 5, 7, and 9 days. (E) Line graph of the MDA content in the MdATG3-overexpressing lines and GL-3 plants under drought conditions at 0, 5, 7, and 9 days. (F) MdMYB16 expression levels in MdATG3-overexpressing lines and GL-3 plants under drought conditions at 0, 5, 7, and 9 days. (G) MdMYB94 expression levels in MdATG3-overexpressing lines and GL-3 plants under drought conditions at 0, 5, 7, and 9 days.

### The overexpression of *MdRLK1* and *MdATG3* increased the level of autophagy in apple leaves

This study quantified autophagosome fluorescence in *MdLRR-RLK1* and *MdATG3* transgenic lines to assess the intensity of autophagy. RT-qPCR analysis revealed significant upregulation of the expression of autophagy-related genes, such as *MdATG3*, *MdATG8C*, and *MdATG8I*, in the *MdLRR-RLK1*-overexpressing lines under salt and drought stress (Fig. 8A-F). As anticipated, we observed a remarkable increase in the number of autophagosomes in the *MdLRR-RLK1* and *MdATG3* overexpression lines under both salt and drought stress compared with that in GL-3 (Supplementary Fig. S5A, B; Fig. 8G, H). Similarly, the expression levels of autophagy-related genes, including *MdATG5*, *MdATG7A*, *MdATG8C*, and *MdATG8I,* were notably increased in the MdATG3-overexpressing lines under salt and drought stress (Fig. 8I-Q). These findings highlight the potential role of enhanced autophagic activity in the improved salt and drought tolerance of *MdLRR-RLK1* and *MdATG3*.

**Fig. 8.**
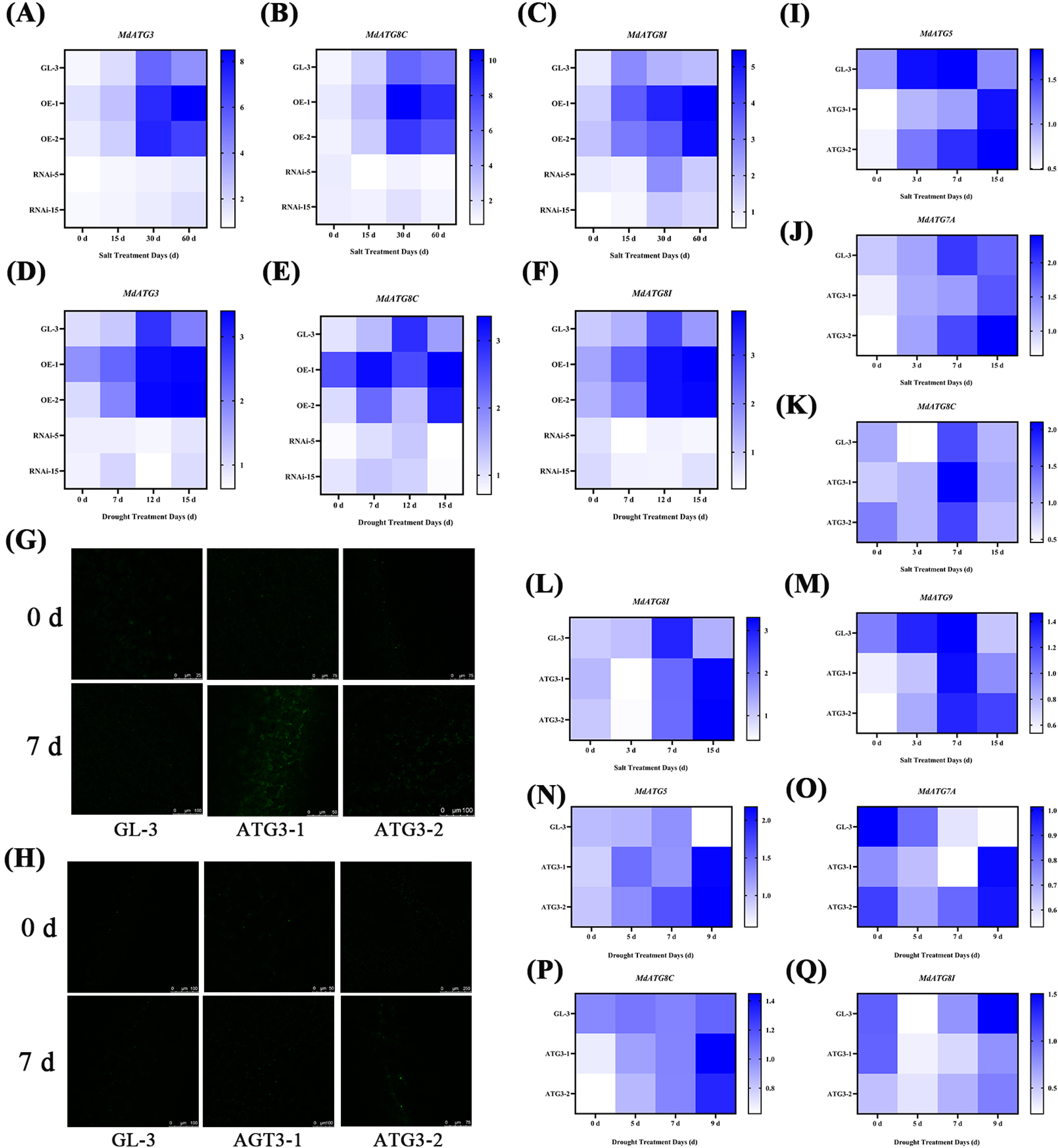
Autophagoso111e fluorescence and RT-qPCR analysis. (A) MdATG3 expression levels in the MdLRR-RLKl transgenic lines and GL-3 plants under salt treat111ent at 0, 15, 30, and 60 days. (B) MdATG8C expression levels in the MdLRR-RLKl transgenic lines and GL-3 plants under salt treat111ent at 0, 15, 30, and 60 days. (C) MdATG8I expression levels in the MdLRR-RLKl transgenic lines and GL-3 plants u11der salt treat111ent at 0, 15, 30, and 60 days. (D) MdATG3 expression levels in the MdLRR-RLKl transgenic lines and GL-3 plants u11der drought conditions at 0, 7, 12, and 15 days. (E) MdATG8C expression levels in the MdLRR-RLKl transgenic lines and GL-3 plants under drought treat111ent at 0, 7, 12, and 15 days. (F) MdATG8I expression levels in the MdLRR-RLKl transgenic lines and GL-3 plants under drought treat111ent at 0, 7, 12, and 15 days. (G) MDC-stained autophagosomes in GL-3 and MdATG3-overexpressing lines under salt stress conditions. The green fluorescent signals represent MDC-stai11ed autophagosomes. (H) MDC-stained autophagoso111es in GL-3 and MdAT.G3-overexpressing lines under drougl1t treat111ent. The green fluorescent signals represent MDC-stained autopl1agoso111es. (I) MdATG5 expression levels in MdAT.G3-overexpressing lines and GL-3 plants under salt stress conditions at 0, 3, 7, and 15 days. (J) MdAT.G7A expression levels i11 the MdATG3-overexpressing lines and GL-3 plants under salt treat111ent at 0, 3, 7, and 15 days. (K) MdATG8C expression levels in the MdATG3-overexpressing lines and GL-3 plants under salt treatme11t at 0, 3, 7, and 15 days. (L) MdATG8I expression levels in the MdAT.G3-overexpressing lines and GL-3 plants under salt treat111ent at 0, 3, 7, and 15 days. (M) MdATG9 expression levels in the MdAT.G3-overexpressing lines and GL-3 plants under salt stress conditions at 0, 3, 7, and 15 days. (N) MdAT.G5 expression levels in MdAT.G3-overexpressing lines and GL-3 pla11ts under drought conditions at 0, 5, 7, and 9 days. (0) MdATG7A expression levels in MdAT.G3-overexpressing lines and GL-3 plants under drought conditions at 0, 3, 7, and 15 days. (P) MdATG8C expression levels in MdAT.G3-overexpressing lines and GL-3 plants under drought conditions at 0, 5, 7, and 9 days. (Q) MdATG8I expression levels in the MdAT.G3-overexpressing lines and GL-3 plants under drought conditions at 0, 5, 7, and 9 days.

### MdATG3 interacts with MdATG8F and MdATG8I-like

Research has identified multiple homologous ATG8 genes in apple. To determine which ATG8 proteins interact with MdATG3, all apple ATG8s were amplified and inserted into vectors for yeast hybrid assays. The results revealed that MdATG3 interacts with MdATG8F (Mdg06B005950) and MdATG8I-like (Mdg09B026360). To confirm this interaction, the *MdATG3*-BT3-N and MdATG8s-PR3-N vectors were used in yeast two-hybrid assays, which revealed that *MdATG3* interacts with MdATG8F and MdATG8I-like in yeast cells (NMY51) (Fig. 9A). BiFC assays further validated this interaction *in vivo*, revealing that MdATG8F/I-like-YFP^n^ and *MdATG3*-YFP^c^ interact in the cell membrane of tobacco, whereas the controls do not produce YFP signals (Fig. 9C). SLCA was also performed by combining MdATG8F/I-like -nLUC and *MdATG3*-cLUC, which resulted in stronger luciferase activity in tobacco leaves than in the negative controls (Fig. 9B). These findings confirm that MdATG8F/I-like can interact with MdATG3 both *in vivo* and *in vitro*.

**Fig. 9.**
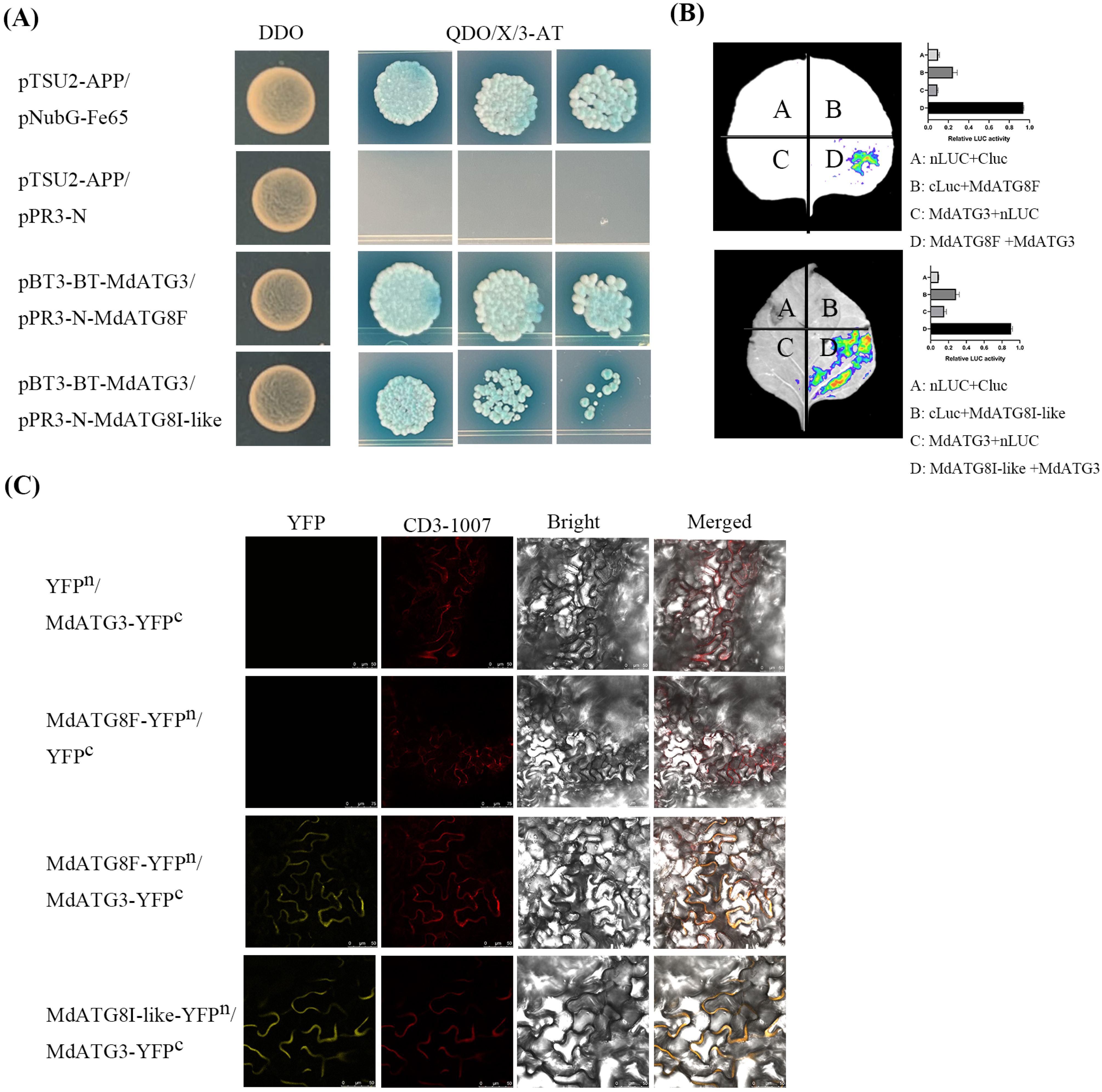
MdATG3 interacts with MdATG8F/I-like in vitro and in vivo. (A) Yeast hybrid validation. The MdATG3-BT3-N, MdATG8F/I-like-PR3-N, and control plasmids were cotransformed into NMY51 yeast competent strains, which were subsequently cultured on media for 3-5 days after transformation. (B) Luciferase imaging assay. MdATG8F/I-like-nLUC, MdATG3-cLUC, and the control plasmid were injected into tobacco leaves via Agrobacterium tumefaciens. What method (reference or reagent kit) should be used to determine LUC content. (C) BiFC experimental analysis. The MdATG8F/I-like-YFPn, MdATG3-YFPc, and control plasmids were subsequently injected into tobacco leaves via Agrobacterium tumefaciens. The control group had scales of 75 µm and 50 µm, whereas the experimental group had a scale of 75 µm.

## Discussion

Apples are a significant fruit variety cultivated in many regions of the world (Bahukhandi et al., 2018), providing essential antioxidants, vitamins, and trace elements to the public (Eberhardt et al., 2000). However, the challenges of drought, land salinization, and climate change threaten apple production (Yamaguchi and Shinozaki, 2006; Fang and Xiong, 2015; Sakuraba et al., 2015; Wei et al., 2017). Therefore, understanding the functions of apple genes and their molecular mechanisms is crucial for breeding highly resistant apple varieties.

### *MdLRR-RLK1* could increase tolerance to abiotic stress

*LRR-RLKs* have gained significant attention in recent years because of their various functions in hormone response, plant stress resistance, growth and development, and signal transduction (Clark et al., 1997; Li and Chory, 1997; Gomez and Boller, 2000; Osakabe et al., 2005; Hord et al., 2006). Furthermore, studies have revealed the nonbiological stress resistance mechanism of *LRR-RLKs*. For example, *AtBAK1* regulates water loss in *Arabidopsis thaliana* by controlling the opening of guard cells (Shang et al., 2016). Similarly, *OsSERK2* not only plays a role in regulating salt tolerance in rice but also participates in the signal transduction of BR (Dong et al., 2020). In Arabidopsis, *AtPEPR1* has been shown to recognize *AtPEP3* and increase tolerance to salt stress (Nakaminami et al., 2018). Additionally, *RLK1* has been identified as a significant player in drought stress (Chen et al., 2021), whereas *AtBRL3* has been found to regulate drought stress tolerance in Arabidopsis through the accumulation of osmoprotectant metabolites such as proline and sugar. In our study, we observed that the overexpression of *MdLRR-RLK1* promoted better growth of apples (Fig. 1), which has also been reported in other species. Among the 223 *LRR-RLKs* identified in Arabidopsis, some have been found to regulate plant stem cell maintenance, organ development, cell expansion, and stomatal development (Nadeau, 2009; Zoulias et al., 2018; Cammarata and Scanlon, 2020; Ou et al., 2021; Zhu et al., 2021). Under drought or salt stress, plants experience disrupted balance, leading to the rapid accumulation of ROS, which can cause membrane lipid peroxidation and damage cell membranes. This results in the leakage of cell contents and, ultimately, cell death. However, plants possess antioxidant enzymes such as SOD, POD, and CAT, which play crucial roles in the clearance of ROS and help maintain plant resistance (Moller, 2001; Bajji et al., 2002; Dievart and Clark, 2004; Cruz and Maria, 2008). Therefore, measuring antioxidant enzyme activity and the MDA content can indirectly indicate plant resistance (Neill et al., 2002; Wang et al., 2010). In our study, we discovered that, compared with the other lines, the *MdLRR-RLK1*-overexpressing lines presented significantly greater antioxidant enzyme activity and lower MDA levels under salt and drought stress. These findings suggest that the increased tolerance of *MdLRR-RLK1* to abiotic stress may be attributed to its strong antioxidant capacity, which is consistent with research findings in peanuts (Wang et al., 2024).

### *MdATG3* could increase tolerance to abiotic stress via autophagy

Autophagy is a well-known process that occurs in all eukaryotic cells and plays a crucial role in the degradation of proteins and organelles (Marshall and Vierstra, 2018). In plants, autophagy remains at a relatively low level during growth and development but is significantly increased in response to stress, aiding in the survival of plants (Wang et al., 2018). Initially, the molecular mechanism of autophagy was studied primarily in yeast and has more recently been expanded to include animals and plants (Tsukada and Ohsumi, 1993; Ohsumi, 2001). The autophagy pathway and the function of ATGs have been validated in both yeast and Arabidopsis, with ATGs playing various roles in autophagy (Kim et al., 2012; Masclaux-Daubresse et al., 2017; Wang et al., 2018; Chung, 2019). Furthermore, autophagy serves as an effective means for plants to cope with salt and drought stress. Silencing *AtATG18a* in Arabidopsis led to increased sensitivity to ROS and the accumulation of peroxides in the transgenic lines (Xiong et al., 2007). Silenced lines of *atg5*, *atg7*, and ATG18a in Arabidopsis also displayed heightened sensitivity to drought and salt stress (Liu et al., 2009; Zhou et al., 2013). On the other hand, the overexpression of ATG18a in apples resulted in increased autophagy activity and increased drought tolerance (Sun et al., 2018). Additionally, evidence suggests that autophagy is involved in the regulation of vacuole blockade of Na^+^ under salt stress (Luo et al., 2017). Recent research has demonstrated that ROS play a role in signal regulation between autophagy and abiotic stress (Chen et al., 2015; Selinski et al., 2018; Zhu et al., 2018). ROS can regulate autophagy by causing damage to plant organelles, subsequently promoting the antioxidant system through the degradation of oxidatively damaged organelles such as mitochondria, chloroplasts, and peroxisomes (Signorelli et al., 2019).

There have been multiple reports discussing the relationship between ROS and autophagy. Xiong et al. (2007) reported that treating Arabidopsis roots with exogenous H_2_O_2_ induced autophagy, resulting in the degradation of oxidative proteins. Furthermore, Pérez-Pérez et al. (2012, 2016) demonstrated that, under stress conditions, plants can modulate autophagy by regulating the oxidation and inactivation of ATG4. Yoshimoto et al. (2009) reported that Arabidopsis *atg2* and *atg5* mutants exhibited excessive accumulation of H_2_O_2_. Yamauchi et al. (2019) reported that the depletion of various ATGs, including ATG2, ATG5, ATG7, ATG10, or ATG12, resulted in reduced antioxidant enzyme activity, thereby compromising plant stress resistance. These findings indicate that autophagy and ROS act synergistically in plants to ensure their ability to withstand adversity. In line with these results, our study revealed that, compared with the control (GL-3), the overexpression of *MdATG3* in apples resulted in increased autophagy ability and increased antioxidant enzyme activity. This phenomenon might explain the strong adaptability of *MdATG3*-1 and *MdATG3*-2 to abiotic stress. During the process of autophagy, ATG8, which functions as an intermediate between E1 and E2, interacts with ATG7 and ATG3, thereby facilitating the connection of ATG8 to the lipid phosphatidylethanolamine. This, in turn, promotes the formation of autophagosomes and enhances plant resistance to abiotic stress (Ichimura et al., 2000; Mallén-Ponce and Pérez-Pérez, 2023). Our study also revealed that MdATG3 interacts with ATG8, further supporting its role in promoting autophagy in apple.

### MdATG3 could regulate autophagy intensity through ubiquitination

Posttranslational modifications, such as phosphorylation, ubiquitination, nitrosylation, and glutathionylation, play significant roles in regulating the state of plants in different environments. Numerous studies have demonstrated the crucial role of ubiquitination in plant immunity (Cheng et al., 2011; Cheng and Li, 2012; Eiyama and Okamoto, 2015; Mach, 2015; Saleh et al., 2015; Xu et al., 2015; Copeland et al., 2016; Gou et al., 2017). The ubiquitination process is an ATP-dependent cascade involving several enzymes, including ubiquitin-activating enzyme E1, ubiquitin-binding enzyme E2, and ubiquitin ligase E3. Typically, E1 activates Ub and transfers it to E2, whereas E3 attaches its specifically recognized substrate to Ub molecules coupled by E2 (Sadanandom et al., 2012; Metzger et al., 2014). Phosphorylation also plays a crucial role (Shen et al., 2023). For example, under carbon starvation, overexpression of AtKIN10 significantly increased the phosphorylation status of ATG1 (Chen et al., 2017). Additionally, DSK2 is phosphorylated by GSK3 to increase plant adaptability to drought and carbon starvation conditions (Nolan et al., 2017).

Yeast hybrid screening revealed an interaction between MdLRR-RLK and MdATG3, which was further confirmed by pull-down, BiFC, LCI, and Y2H assays. Although ATG3-1 and ATG3-2 presented greater resistance to salt and drought stress than did GL-3, their resistance was not as strong as that of the overexpression lines OE-1 and OE-2 of MdLRR-RLK1. Autophagic fluorescence in the OE lines was not weaker than that in the ATG3 lines, but the opposite was observed in the RNAi-5 and RNAi-15 lines. These results suggest that the abundance of MdLRR-RLK1 may play a significant role. MdLRR-RLK1 functions as a kinase but does not phosphorylate ATG3, while ATG3, an E2 ubiquitin-conjugating enzyme, targets phosphorylated LRR-RLK1 for degradation, reducing its abundance and enhancing autophagy. This could explain why OE-1 and OE-2 presented stronger abiotic stress resistance than did ATG3-1 and ATG3-2.

In summary, the relationship between MdLRR-RLK1 and MdATG3 has been established (Fig. 10), and mechanisms regulating apple growth and stress resistance have been proposed. These findings provide a strong foundation for future studies on the molecular mechanisms and development of salt- and drought-resistant apple varieties.

**Fig. 10.**
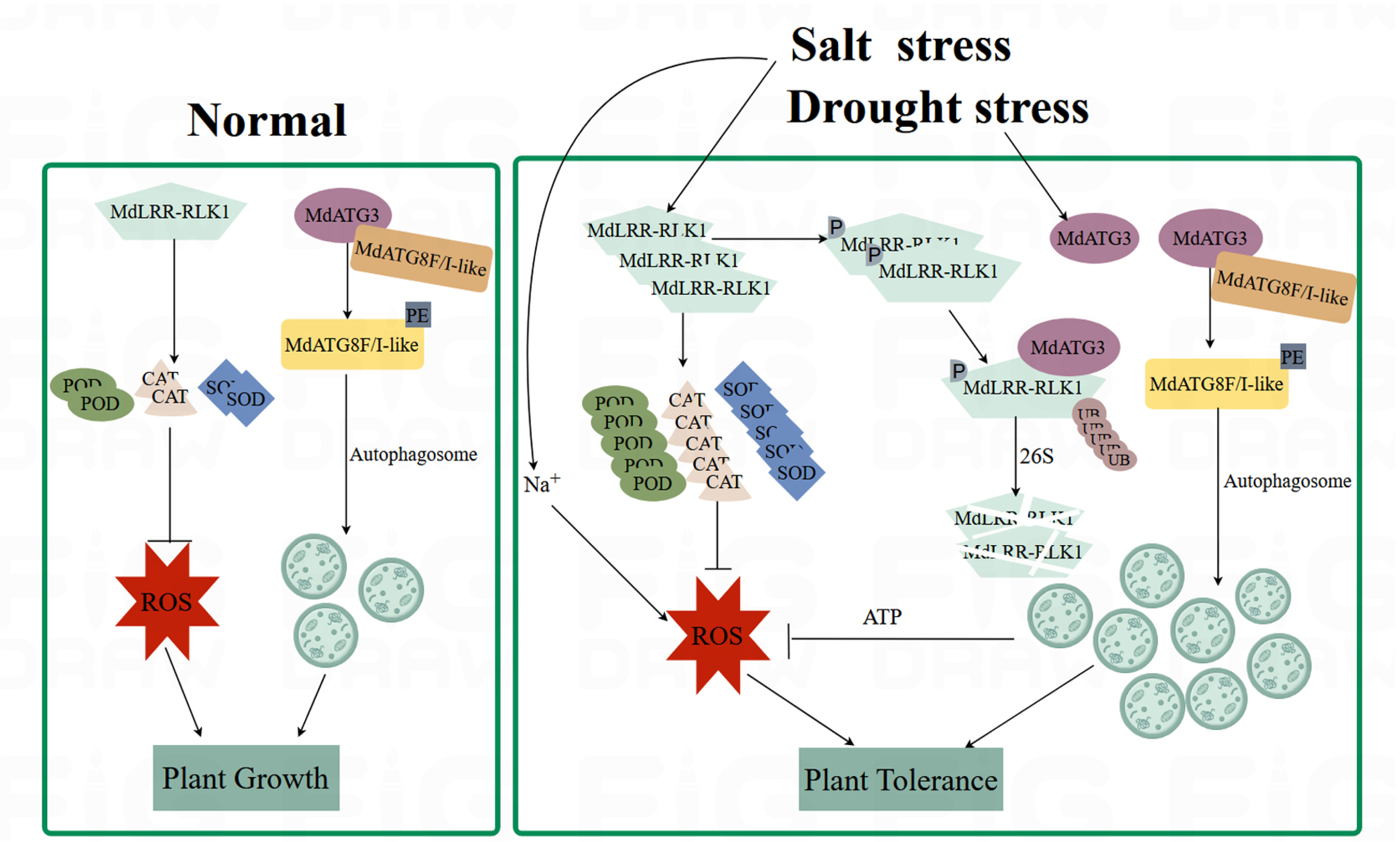
Illustrates the proposed mechanism of MdLRR-RLKl and MdATG3 in the abiotic stress tolerance of apple. Under normal conditions, MdLRR-RLKl promotes growth in terms of content, whereas MdATG3 interacts with MdATG8F/I-like to promote growth through autophagy. Under salt or drought stress, both MdLRR-RLKl and MdATG3 are induced. MdLRR-RLKl increases antioxidant enzyme production, reduces ROS and enhances stress tolerance. Concurrently, MdATG3 enhances autophagy via acetylation of MdATG8F/I-like, further improving stress tolerance. MdATG3 also degrades phosphorylated MdLRR-RLKl through the 26S proteasome, reducing ROS levels and contributing to stress tolerance.

## Materials and methods

### Plant materials and growth conditions

This study used wild-type apple plants (*Malus domestica* Borkh) ‘GL-3’. The GL-3 tissue culture-generated seedlings were initially cultured in a subculture medium called Phytotech Labs HCE0519387A. Before transplantation, the seedlings were cultivated in rooting media for three months to ensure root system development (Coolaber Science and Technology PM322028700). During transplantation, both rooted GL-3 and transgenic lines were placed in a substrate composed of peat, vermiculite, and perlite at a ratio of 3:2:1. The substrate was covered with film to maintain moisture. After 20 days of cultivation under the film, the film was removed, and cultivation continued for an additional two to five months. The plants were cultivated at a temperature of 25°C under a photoperiod of 12 hours of light and 12 hours of darkness (Dai et al., 2013).

### Stable genetic transformation

To achieve stable genetic transformation, the full-length CDS of *MdLRR-RLK1* was inserted into the pRI-101-AN vector. The constructed vector was subsequently transformed into *Agrobacterium tumefaciens* strain EHA105. The GL-3 tissue-cultured seedlings were then used for stable genetic transformation following the method described by Liu et al. (2024).

### qRT‒PCR

Total RNA was extracted via the CTAB method as described by Chang et al. (2007). cDNA synthesis was performed via the PrimeScript RT reagent Kit with gDNA Eraser (Takara RR047A). Real-time quantitative PCR (RT‒qPCR) was carried out via SYBR Premix Ex Taq (Cwbio UltraSYBR Mixture CW2601H) on an IQ6 Real-Time PCR instrument (Applied Biosystems QuantStudio 6 FLEX). Actin was used as the internal control for apple, and data analysis was conducted via the 2^-ΔΔCt^ method (Livak and Schmittgen, 2001). The RT-qPCR primers used in this study are detailed in Table S1.

### Salt stress treatment

The apple plants were subjected to a 60 days treatment with 200 mM NaCl to observe the phenotype in response to salt stress. Photos were taken, and samples were collected at intervals of 0, 15, 30, and 60 days for subsequent experiments.

### Drought stress treatment

Each apple plant used for the drought treatment was irrigated with 50 mL of water during cultivation. Watering was stopped when the treatment started. Photos were taken, and samples were collected at 0, 3, 5, and 7 days for subsequent experiments.

### Enzyme activity assays

A total of 0.1 g of sample was weighed, and the enzyme mixture was extracted via 0.05 mol/L PBS buffer at 4°C for 3 days. The extracted solution was then centrifuged at 4°C and 12,000 rpm for 20 minutes. The supernatant obtained after centrifugation was used as the enzyme mixture. The determination of peroxidase (POD), catalase (CAT), and superoxide dismutase (SOD) activity was based on Mahmood et al. (2016).

### MDA assay

The method for determining the MDA content was the protocol developed by Cakmak and Horst (1991), with minor adaptations. Leaf and root samples weighing 1.0 g each were pulverized in a 0.1% (w/v) trichloroacetic acid (TCA) solution. The homogenate was subsequently centrifuged at 12,000 rpm for 15 minutes. To the supernatant, 0.5% thiobarbituric acid (TBA) was added. The mixture was heated in a 95°C water bath for 50 minutes and then rapidly cooled in an ice bath to halt the reaction. Next, the sample was centrifuged again at 12,000 rpm for 10 minutes, and the absorbance of the supernatant was measured at 450 nm, 532 nm, and 600 nm. The quantification of MDA was performed via the spectrophotometric method on the basis of its molar absorption coefficient (155 mM⁻¹ cm⁻¹).

### Subcellular localization

The subcellular localization method followed the protocol of Wang et al. (2019). The coding sequences of *MdLRR-RLK1* and *MdATG3* without stop codons were subsequently cloned and inserted into the vectors *MdLRR-RLK1*-GFP and *MdATG3*-GFP, respectively. *Nicotiana benthamiana* leaves were then infiltrated with *Agrobacterium tumefaciens* strain GV3101. Following a 48-hour dark incubation and a 24-hour normal light incubation, GFP fluorescence was visualized via laser confocal microscopy (TCS SP8, Leica, Germany, with a scanning resolution of 8192 * 8192 pixels). CD3-1007 mCherry serves as a cell membrane marker (Wang et al., 2019).

### Yeast library screening and validation

GL-3 tissue culture-generated seedlings were cultivated for a period of 3 months. Samples were taken from the leaves and roots to construct a yeast nuclear membrane dual protein library. The library was developed by Shanghai OE Biotech Co., Ltd. (https://www.oebiotech.com/). The full-length CDS of *MdLRR-RLK1* was introduced into the vector *MdLRR-RLK1*-BT3-STE and used for protein screening. The screening library method was provided by Shanghai OE Biotech Co., Ltd. The screening was performed via SD/-Trp/-Leu/-His/-Ade media supplemented with 40 μg/mL X-α-gal and 30 mM 3-AT (QDO/X/3-AT). Yeasts obtained from the screening were sequenced via Sanger sequencing. After the corresponding vector was constructed, the screened genes were validated in QDO/X/3-AT for a period of 3-5 days at 30°C.

### Bimolecular fluorescence complementation assay (BiFC assay)

The bimolecular fluorescence complementation (BiFC) assay followed the protocol established by Walter et al. (2004). The full-length coding sequences of *MdLRR-RLK1* and *MdATG3* were subsequently cloned and inserted into the vectors *MdLRR-RLK1*-YFP^n^ and *MdATG3*-YFP^c^, respectively. The subsequent steps and the use of a cell membrane marker were in accordance with the subcellular localization method.

### LCA assay

The LCA method was adapted from Zhou et al. (2018). The full-length coding sequences of *MdLRR-RLK1* and *MdATG3* were subsequently cloned and inserted into the vectors *MdLRR-RLK1*-nLuc and *MdATG3*-cLuc, respectively. The subsequent steps were consistent with the subcellular localization method. Fluorescence measurements were taken via the Night SHADE evo *In Vivo* Plant Imaging System (LB 985 N, Berthold, Germany).

### Pull-down assay

The pull-down assay followed the protocol described by Li et al. (2024). The full-length coding sequences of *MdLRR-RLK1* and *MdATG3* were subsequently cloned and inserted into the vectors *MdLRR-RLK1*-His and *MdATG3*-GST, respectively. These constructs were subsequently transformed into competent *Escherichia coli* BL21 (DE3) cells, which were subsequently induced with IPTG. The bacterial cells were harvested, and the proteins were extracted via Tiachi *E. coli* Lysis Buffer (ACE Biotechnology, BR0005-2). Protein purification was performed via Ni NTA Resin (TransGenBiotech, DP101-01) for *MdLRR-RLK1*-His and GST Resin (TransGenBiotech, DP201-01) for *MdATG3*-GST. The antibodies used included mouse anti-His-Tag monoclonal antibody (ABclonal, AE003), mouse anti-GST-Tag monoclonal antibody (ABclonal, AE001), and HRP-conjugated goat anti-mouse IgG (H+L) (ABclonal, AS003).

### Bioinformatics analysis

TAIR (https://www.arabidopsis.org/) was used to search for genes related to LRR-RLKs in Arabidopsis. MEGA7 software was used to construct a phylogenetic tree. Pfam (https://pfam.xfam.org/) and SMART (http://smart.embl-heidelberg.de/) were used to perform protein domain analysis. DNAMAN version 9 software was used for visualizing and annotating the relevant domains.

### MDC staining

After treatment, the apple leaves were cut into small fragments and fully immersed in 100 μM MDC staining solution. Vacuum infiltration was used to ensure that the staining solution penetrated the entire leaf tissue. After being stained for 3-4 hours, the leaves were washed twice with 1x PBS buffer. Subsequent imaging and observation were carried out via a confocal laser scanning microscope. The MDC fluorescence channel was excited at a wavelength of 405 nm, and the emitted light was captured between 410 and 585 nm.

### Phosphorylation and ubiquitination experiments

The *in vitro* phosphorylation assay method was adapted from Li et al. (2023). The antibodies used were a pan phospho-serine/threonine rabbit polyclonal antibody (Beyotime, AF5725) and HRP-labeled goat anti-rabbit IgG (H+L) (Beyotime, A0208). The recombinant proteins *MdLRR-RLK1*-His and *MdATG3*-GST were overexpressed, and phosphatase inhibitors (phosphate inhibitor cocktail A from Beyotime, product number P1081) were added during bacterial lysis to obtain proteins for Western blot (WB) analysis. The LC-MS/MS analyzed by Shanghai Applied Protein Technology Co., LTD.

Using *MdLRR-RLK1*-His as the kinase and *MdATG3*-GST as the substrate, the kinase and substrate were mixed in a 1:5 ratio with kinase assay buffer (25 mM Tris-HCl [pH 7.5], 10 mM MgCl_2_, 1 mM CaCl_2_, 10 mM ATP, and 1 mM DTT) and incubated at 30°C for 2 hours. The reaction was then stopped by heating with 1x SDS loading buffer at 100°C for 5 minutes. The protein mixture was separated on a 12% SDS-PAGE gel. WB detection was performed using an anti-phospho serine/threonine antibody (Beyotime, AF5725).

The *in vitro* ubiquitination detection reaction system consisted of 50 μL, containing 25 mM Tris-HCl [pH 7.5], 100 mM NaCl, 5 mM MgCl_2_, 5 mM ATP, 5 μg of ubiquitin, 150 nM E1 (Proteintech Group Inc, Ag8920), 150 nM E3 (Proteintech Group Inc, Ag0346), 5 ng of purified *MdLRR-RLK1*-His, and 5 ng of purified *MdATG3*-GST proteins. The mixture was incubated at 30°C for 2 hours and then stopped by heating with 1x SDS loading buffer at 100°C for 5 minutes. The protein mixture was separated on a 12% SDS-PAGE gel. WB detection was performed using anti-ubiquitin (Ub) (Bioss, bsm-60167R).

## Acknowledgements

We would like to thank Dr Huixia Shou at Zhejiang University for providing the CD3-1007 vector.

## Author contributions

HY Dai designed research. WJ Chen, W Guo, C Zhang and Y Zhao performed research., WJ Chen, YY Lei, C Chen and ZW Wei analyzed data. WJ Chen drafted the manuscript. W Guo and HY Dai revised the manuscript.

## Supplementary data

The following materials are available in the online version of this article.

Supplementary Fig. S1. Analysis of the genetic characteristics of MdLRR-RLK1.

Supplementary Fig. S2. Transcriptomic data analysis of the *MdLRR-RLK1* transgenic lines.

Supplementary Fig. S3. Phosphorylation of MdLRR-RLK1 and MdATG3 protein.

Supplementary Fig. S4. Gene analysis and phenotypic characteristics of *MdATG3*.

Supplementary Fig. S5. MDC-stained autophagosomes in GL-3 and *MdLRR-RLK1* transgenic lines under salt stress conditions.

Supplementary Table. S1. The primers used in this study.

## Data availability

The authors confirm that the data supporting the findings of this study are available within the article and its Supporting Information.

